# Establishment of a patient-derived adrenocortical carcinoma 3D tumor construct platform for evaluation of therapeutic strategies

**DOI:** 10.64898/2026.01.08.698468

**Authors:** Marco A. Rodriguez, Hemamylammal Sivakumar, Liudmila Popova, Saeed Derakhshesh, Daniel Chopyk, Wayne O. Miles, Priya H. Dedhia, Aleksander Skardal

## Abstract

Adrenocortical carcinoma (ACC) is an under-studied, aggressive cancer of the adrenal glands where surgical resection is currently the only effective curative option. However, after surgery the majority of patients experience tumor progression. The median overall survival of 12 months has not improved since approval of mitotane in 1970. The lack of effective therapies in ACC is partially due to the lack of preclinical models that accurately represent human ACC. Most attempts to generate ACC cell lines or animal models have been unsuccessful. The few existing models do not adequately reflect the oncogenic signaling pathways or intratumoral heterogeneity of human ACC. These limited model systems have hindered identification of drivers of tumor progression and immune escape mechanisms, thereby limiting development and testing of novel therapeutic approaches such as targeted therapies or immunotherapies. We developed human ACC patient-derived tumor constructs (PTCs) encapsulated in synthetic extracellular matrix. Our ACC PTCs exhibit hallmarks of ACC: proliferation, expression of key ACC biomarkers, such as SF1, and production of cortisol. We provide characterization in the form of immunofluorescence staining for ACC biomarkers, and confirmation of PTC proliferation and cortisol production – a hallmark of ACC. We then demonstrate the utility of ACC PTCs for evaluation of chemotherapies currently used clinically (mitotane) with and without experimental cocktails of etoposide, doxorubicin, and cisplatin, experimental targeted therapies, and cellular immunotherapies, primarily in the form of natural killer (NK) cell therapy. In particular, the latter – cellular immunotherapies – are cutting edge studies demonstrating potential to evaluate immunotherapies in ACC clinical scenarios. Together these data provide evidence that patient-derived ACC models can serve as an important tool for identification of future points of intervention and testing of novel therapeutic strategies to improve ACC clinical care.

## 1. Introduction

Adrenocortical carcinoma (ACC) is an aggressive and understudied cancer of the adrenal glands with an annual incidence of 200-400 cases in the United States.^1,2^ Drug treatments have been largely ineffectual, making surgical resection is the only curative option. Unfortunately, the majority of patients present with unresectable disease or experience tumor recurrence after surgical intervention.^3^ The 5-year progression-free survival rate for metastatic ACC is 6%.^4^ Despite this prognosis, the median survival of 12 months has not improved since approval of mitotane in 1970.^3^ Current first-line treatment with mitotane with or without etoposide, doxorubicin, and cisplatin (EDP) extends median progression-free survival by just 5 months without improving overall survival, underscoring the urgent need for new therapies.^5^

The lack of therapeutic progress in ACC is partially attributable to the scarcity of preclinical models that accurately represent human ACC. Most attempts to generate ACC cell lines or animal models have been unsuccessful. The few existing models fail to adequately reflect the oncogenic signaling pathways or intratumoral heterogeneity of human ACC. These constrained model systems have hindered identification of drivers of tumor progression and immune escape mechanisms, thereby limiting development and testing of novel therapeutic approaches.^3,4^ While patient-derived ACC organoids have been utilized in select studies, they, like many patient-derived tumor models – rely on Matrigel, a murine sarcoma-derived biomaterial that introduces unwanted variables and significant lot-to-lot variability that can confound experimental results.^6^ An ACC model system that accurately recapitulates human disease would provide a critical platform for testing novel targeted therapies and immunotherapeutic approaches. Many ACC tumors produce excess cortisol that suppresses immune cell activity, creating a significant barrier to therapeutic efficacy.^7^ Physiologically relevant models are therefore essential to develop strategies that can overcome this challenge.

To address the limitations of existing tumor models, our group has developed patient-derived tumor constructs (PTCs) – 3D tumor generated from patient biospecimens that can incorporate immune system components. Over the past decade, we have successfully generated PTCs from multiple cancer types, including colorectal, appendiceal, lung adenocarcinoma, mesothelioma, melanoma, and glioma tumors and deployed these models for both chemotherapy and immunotherapy screening studies.^8–18^ Here, we describe the first 3D bioengineered tumor models from clinical ACC biospecimens using a defined and bioengineered extracellular matrix (ECM) hydrogel system. These ACC PTCs maintain classical ACC biomarker expression, cortisol production, and proliferation over time in culture. We first validate the platform using standard-of-care mitotane and EDP chemotherapy then apply targeted pathway inhibitors and natural killer cell-based immunotherapy to assess several novel therapeutic interventions.

## 2. Results

### 2.1. Overall study design

To preserve intratumoral heterogeneity present in patient malignancies, we generated ACC PTCs by incorporating cells from clinical biospecimens into functionalized collagen and hyaluronic acid hydrogels (**Figure 1**). In this 3D platform, we characterized cell proliferation, expression of typical ACC markers, and production of cortisol, a hormone secreted in excess by up to 60% of ACC patients.^19^ PTCs were then evaluated for response to current chemotherapeutics, targeted therapies, and cellular immunotherapy. While the ACC cell line, NCI-H295R, cell line has been extensively implemented in multiple cancer models, including our team’s metastasis platform, these cell line-based systems lack the intratumoral heterogeneity present in patient tumors.^20,21^ Our PTC approach addresses this limitation.

**Figure 1.**
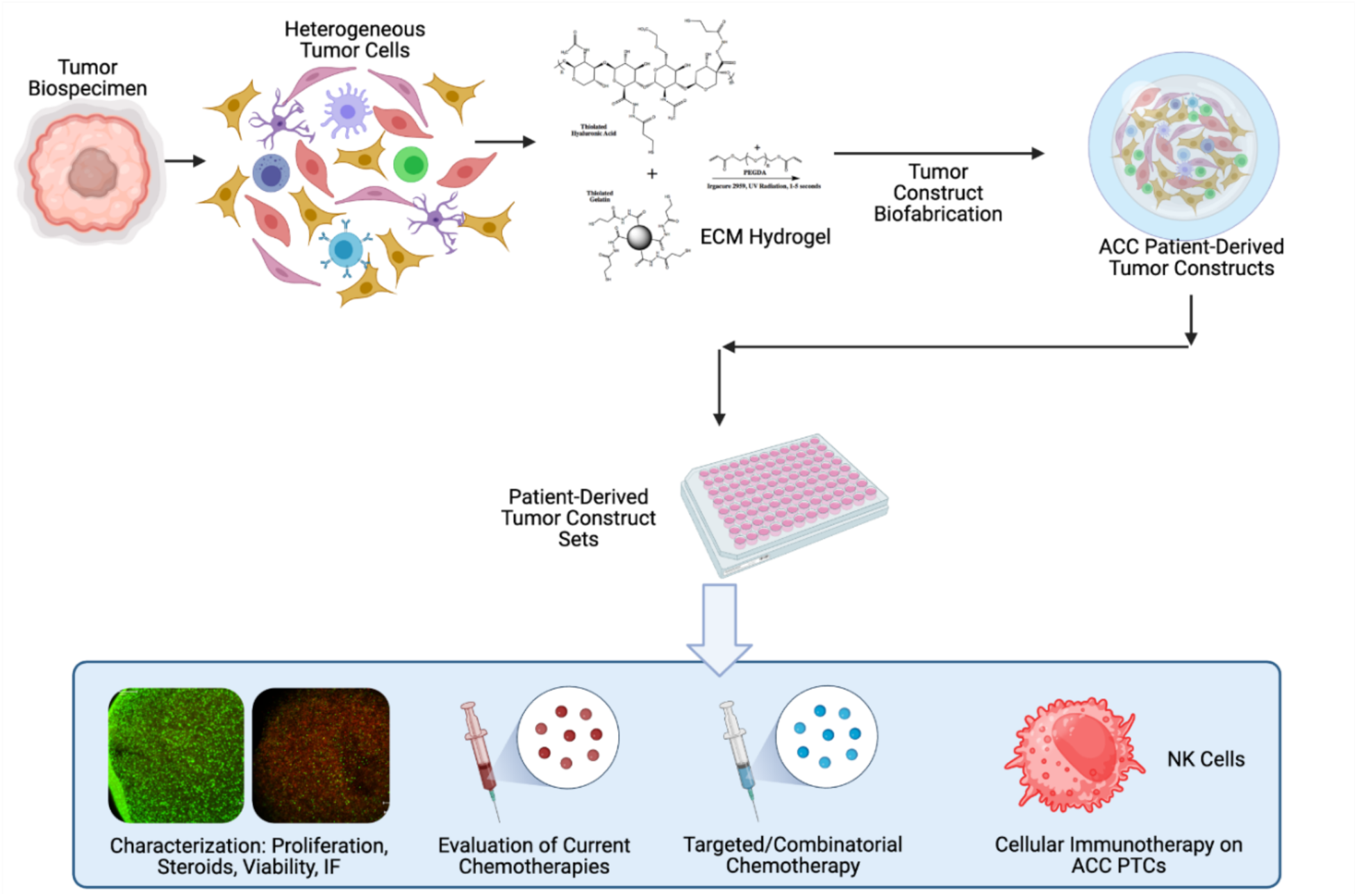
*ACC* Experimental design overview. ACC tumor biospecimens were dissociated and encapsulated in a hyaluronic acid-based hydrogel to form ACC PTCs which were subjected to characterization studies, drug screens using current chemotherapies and targeted/ combinatorial chemotherapy intervention, and evaluation of cellular immunotherapies.

### 2.2. Phenotypic and functional characterization

PTCs were characterized by immunofluorescence staining for proteins commonly expressed or dysregulated in ACC to confirm maintenance of tumor identity. IGF2 and nuclear β-catenin, key drivers of ACC tumorigenesis through dysregulation of insulin-like growth factor and WNT signaling pathways – which are upregulated in most ACC tumors – ^22^ were expressed throughout most PTC cells (**Figure 2 a, c, e**). SF-1, a transcription factor essential for adrenocortical development and steroidogenesis, and alpha-inhibin and Melan-A, markers of adrenocortical lineage, were also expressed, and are key markers used to diagnose ACC pathology (**Figure 2 b, d**).^23,24^

**Figure 2.**
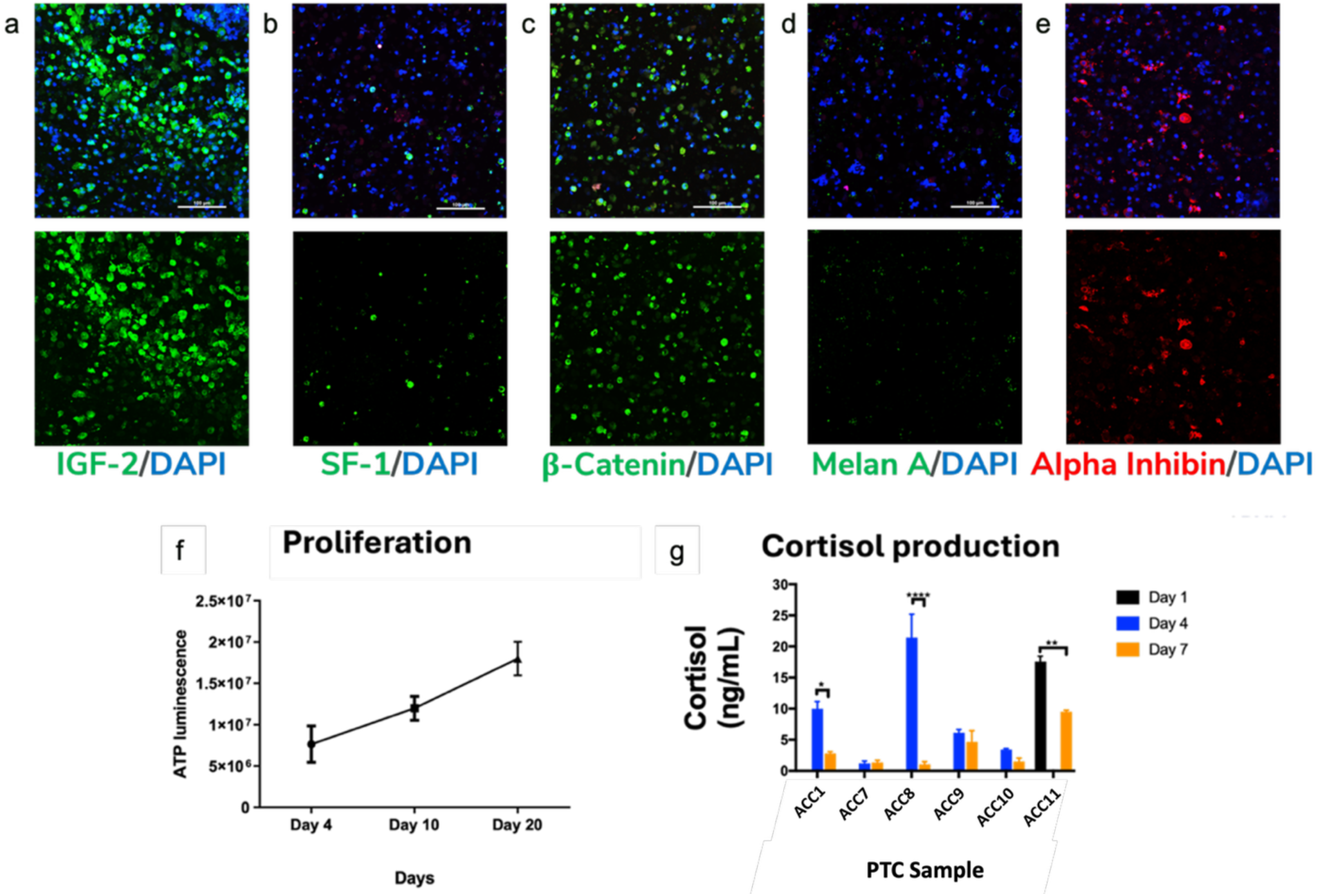
ACC PTCs maintain expression of ACC biomarkers, proliferate, and produce cortisol. a-e) Immunofluorescence staining of a) IGF-2, b) SF-1, c) ß-catenin, d) melan A, and e) alpha inhibin. Top panels: Green or Red – Indicated protein; Blue – DAPI counterstain. Bottom panels: Green or Red – Indicated protein only. f) Representative measurement of ATP at days 4, 10, and 20 in PTCs, showing proliferation over time. g) Measurement of secreted cortisol in PTC cultures for 6 PTC sets on days 4 and 7 (or 1 and 7 in one case). Statistical significance: * p<0.05; ** p<0.01; **** p<0.001.

We assessed ACC PTC viability by measuring ATP activity. PTCs demonstrated sustained proliferation over a 20-day period, with measurements on days 4, 10, and 20 (**Figure 2 f**). Sustained viability at day 20 confirmed the compatibility of the collagen and hyaluronic acid 3D system for ACC PTCs.

Cortisol production is a hallmark feature of ACC and contributes to dysregulation of multiple biological processes. We measured cortisol secretion by ELISA on days 1, 4, and 7. Production levels varied across patient samples but typically peaked at day 4 in culture (**Figure 2 g**). Most PTC sets exhibited declining cortisol levels over time, indicating that shorter culture periods (≤7 days) are optimal for cortisol-related studies in this model.

### 2.3. Transcriptomic characterization of PTCs

Despite wide usage, The NCI-H295R cell line, does not recapitulate inter-patient tumor heterogeneity. We hypothesized that ACC PTCs preserve the transcriptomic profiles of their tumors of origin. To test this hypothesis, we performed bulk RNA sequencing on four PTC-tumor pairs alongside NCI-H295R cells. Principal component analysis and hierarchical clustering demonstrated that PTCs clustered more closely with their corresponding tumors than with NCI-H295R (**Figure 3a-b**), confirming that PTCs faithfully preserve patient-specific transcriptional signatures—a key advantage over immortalized cell lines.

**Figure 3.**
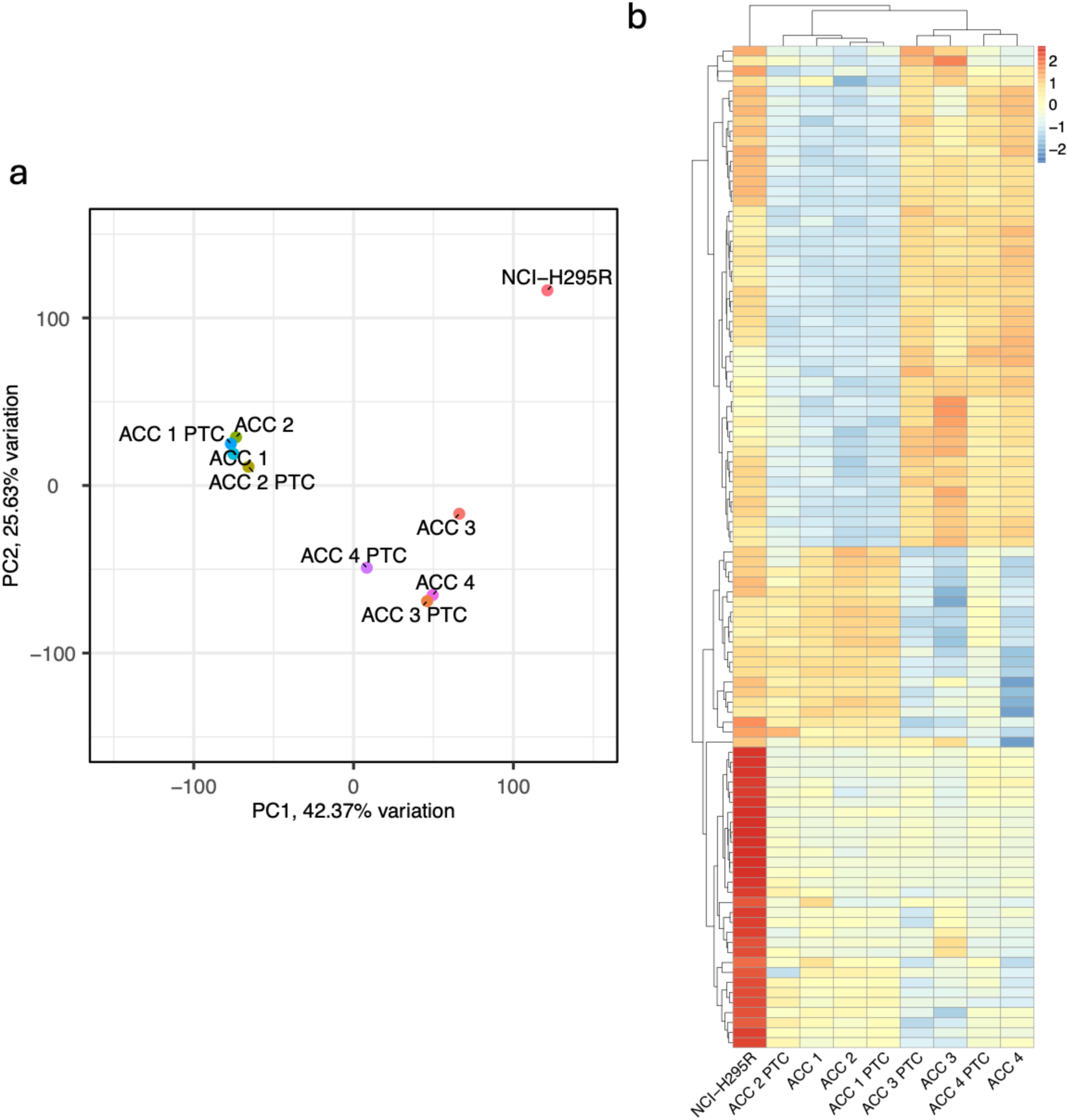
ACC PTCs maintain patient-specific transcriptional signatures. A) Principal component analysis and B) heatmap of genes driving principal component separation across PTCs, matching tumors, and NCI-H295R cells, demonstrating that PTCs cluster more closely with their corresponding tumors than with NCI-H295R cells.

### 2.4. Drug screening with current clinical therapeutics

We evaluated current ACC treatment regimens, mitotane monotherapy and mitotane plus EDP,^25^ —by treating PTCs with varying EDP concentrations combined with 8 μg/mL mitotane (**Figure 4**). Cell viability was assessed by ATP quantification and LIVE/DEAD staining when cell yield was sufficient (**Figure 4a-d**) or by ATP quantification alone when limited(**Figure 4e-f**). Mitotane monotherapy demonstrated minimal cytotoxic effects across all PTC sets. Higher EDP concentrations, particularly in combination with mitotane, reduced viability and ATP activity, though these differences did not consistently reach statistical significance. LIVE/DEAD imaging confirmed these findings, revealing increased ethidium homodimer-1 signal in combination-treated cells. Metastatic (hepatic) PTCs showed greater sensitivity to EDP alone and mitotane plus EDP than paired primary PTCs from the same patient (**Figure 4f**), though differences did not reach statistical significance.

**Figure 4.**
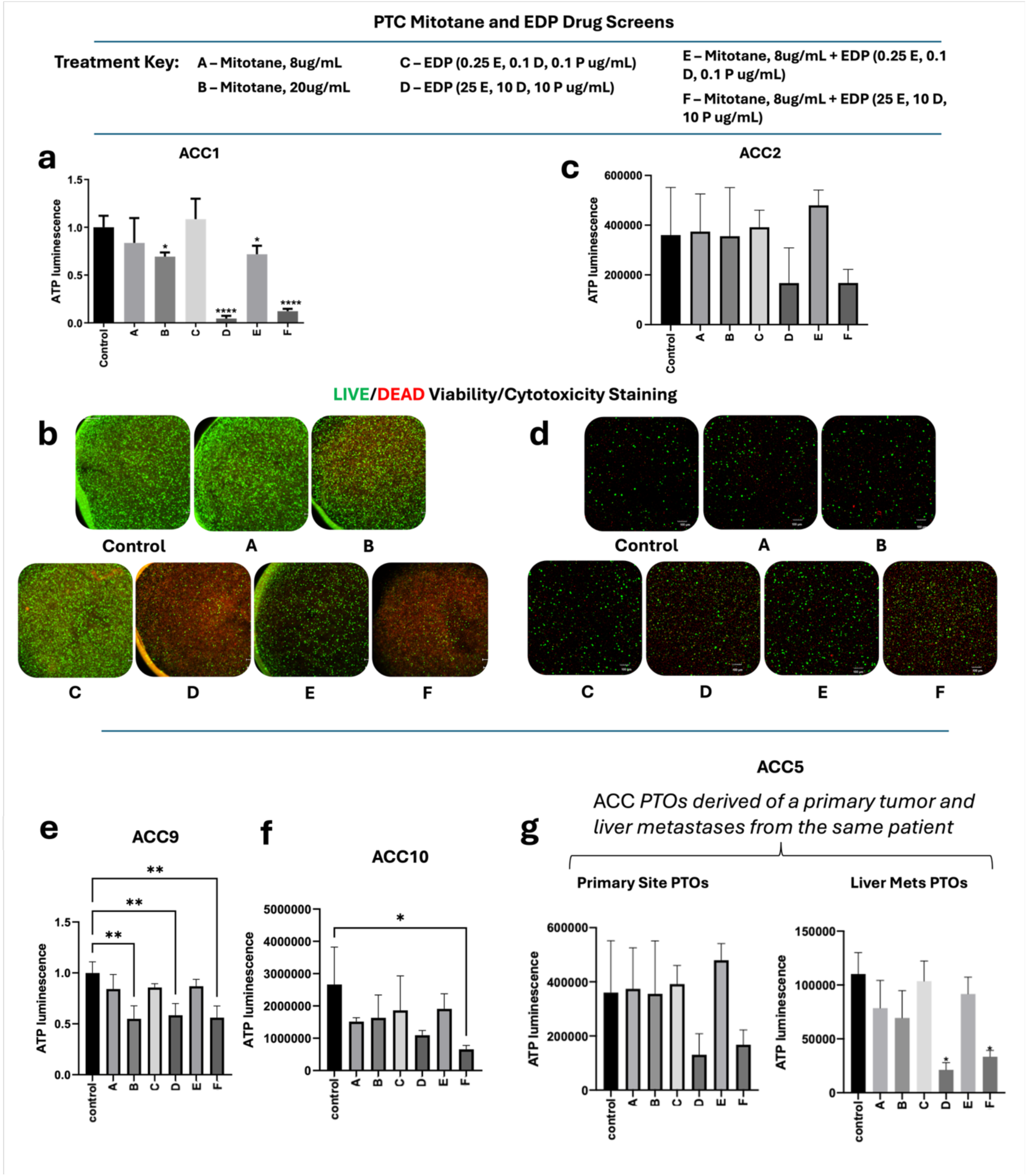
Mitotane and EDP studies show variable responses between PTC sets. a-b) ATP and LIVE/DEAD drug response data for PTC set 101. c-d) ATP and LIVE/DEAD drug response data for PTC set 1123. e-f) ATP drug response data for PTC sets 503 and 531; g) ATP drug response data for PTCs derived from a primary ACC tumor and a metastatic ACC tumor from the same patient. Drug concentrations/combinations are described in the included key. Statistical significance: * p<0.05; ** p<0.01; **** p<0.001. LIVE/DEAD staining: Green – Calcein AM-stained viable cells; Red – Ethidium homodimer-1-stained dead cell nuclei.

**Figure 5:**
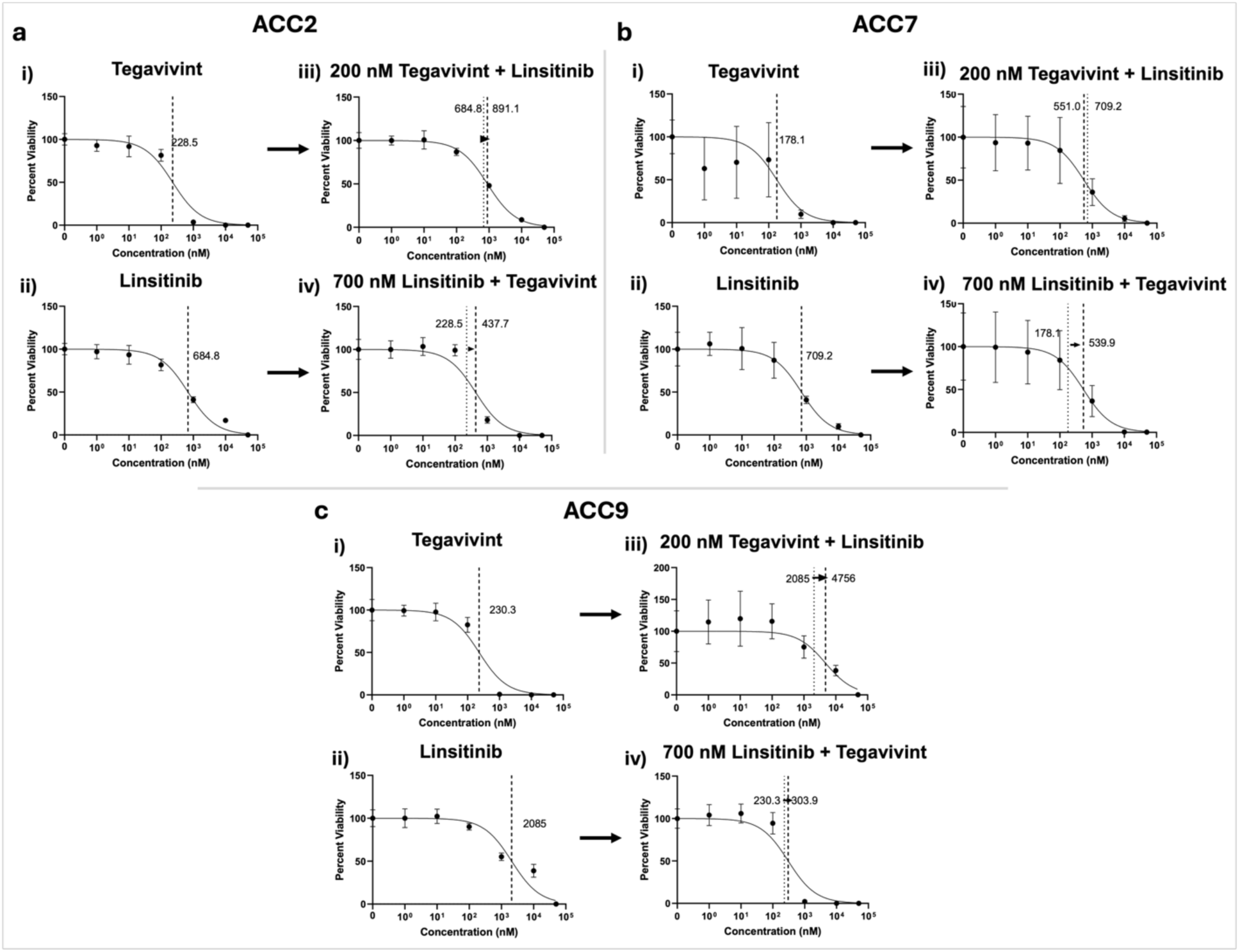
Monotherapy with tegavivint and linsitinib is as effective or superior to combinatorial treatment in ACC PTCs. a-c) Dose response curves for tegavivint and linsitinib treatments of ACC PTCs for a) ACC2, b) ACC7, and c) ACC9. Each set of PTCs was treated with i) tegavivint or ii) linsitinib alone initially to identify approximate IC50 values. Subsequently, combinatorial treatments were tested. Each set of PTCs was treated with iii) 200 nM tegavivint with varying linsitinib concentrations, and iv) 700 nM linisitinib with varying tegavivint concentrations. Arrows between vertical dashed lines show changes in IC50 due to combinational treatments.

### 2.5. Inhibition of Wnt, IGF2, P53, and ATR pathways

The most aggressive ACC tumors exhibit dysregulation of IGF2, Wnt/β-catenin, and cell cycle pathways.^26,27^ Based on these pathway alterations, we evaluated targeted inhibitors of Wnt signaling (tegavivint and XAV-939), IGF2 signaling (linsitinib), and TP53 (NSC59984). We also tested ATR inhibitors (VX-803 and elimusertib), as we recently identified ataxia telangiectasia-mutated and Rad3-related (ATR) inhibition as a promising therapeutic strategy for ACC.^28^

We first screened these compounds in NCI-H295R tumor constructs to establish dose-response curves across multiple concentrations. Cell viability was assessed by ATP quantification and LIVE/DEAD staining to determine IC50 values (**Supplemental Figures 1-7**). ATR inhibitors demonstrated the lowest effective concentrations – 4.5 and 39 nM for VX-803 and elimusertib, respectively, followed by the Wnt inhibitor tegavivint with an IC50 of 214.7 nM. XAV-939 showed minimal response even at high concentrations. While linsitinib and NSC59984 reduced tumor viability, both required substantially higher concentrations to achieve cytotoxic effects at IC50 values of 1.3 and 2.2 μM, respectively (**Supplemental Figure 1**).

Given limited patient biospecimen availability, we selected two compounds for ACC PTC testing: linsitinib (IGF2 inhibitor) and tegavivint (Wnt inhibitor), based on established IGF2 and Wnt pathway dysregulation in ACC. Initially, we evaluated whether these compounds demonstrated synergistic effects in NCI-H295R tumor constructs. Combination treatment in NCI-H295R cells showed additive cytotoxicity in some conditions (**Supplemental Figure 1e-i**) but not others (**Supplemental Figure 1e-ii**). Specifically, when we screened a range of tegavivint concentrations paired with 1.3 μM linsitinib treatment, we observed a decrease in the IC50 value by a modest 24 nM. Alternatively, when we screened a range of linsitinib concentrations paired with 200 nM tegavivint treatment, we observed a negligible decrease in the IC50 value.

Ultimately, these data provided motivation to test these drugs independently and in combination in ACC PTCs. Both tegavivint and linsitinibmonotherapy significantly reduced viability in ACC PTCs across three patient samples in a dose-dependent manner with IC50 values of approximately 200 nM and 700 nM, respectively, showing some consistency between PTC sets (**Figure 4a-ci**). However, combination treatment was not superior to monotherapy (**Figure 4a-cii**). In contrast to the effects observed in NCI-H295R cells, combination treatment in ACC PTCs unexpectedly resulted in higher IC50 values in some cases, suggesting antagonistic interactions.

### 2.6. Evaluation of cellular immunotherapy

Beyond small molecule inhibitors, we evaluated cellular immunotherapy as an alternative therapeutic approach given the growing clinical interest in natural killer (NK) and T cell-based therapies for solid tumors. As proof of concept, we deployed the NCI-H295R ACC cell line in a tumor immune construct (TIC) consisting of NK and T cell lines to assess cytotoxic function. Untreated controls maintained CFSE-stained tumor cells with minimal SYTOX Blue signal, indicating low cell death indicating minimal cell death (**Supplemental Figure 8**). Introduction of NK-92 cells decreased CFSE signal (tumor cells), increased PKH26 signal (infiltrated NK-92 cells), and elevated SYTOX Blue (cell death), suggesting successful NK cell infiltration and effector function (**Supplemental Figure 8**).(**Supplemental Figure 8**). In contrast, TALL-104 T cells showed immunofluorescence patterns similar to untreated controls, indicating ineffective cytotoxicity in this model (**Supplemental Figure 8**).

Based on NK-92 efficacy in NCI-H295R constructs, we evaluated this approach in two ACC PTC sets using the same experimental workflow. With introduction of NK cells, both PTC sets demonstrated decreased CFSE fluorescence (labeled tumor cells), increased PKH26 fluorescence (infiltrated NK-92 cells), and increased SytoxBlue (cell death) (**Figure 6a-b**), confirming NK-92 cytotoxicity in patient-derived ACC models.

**Figure 6.**
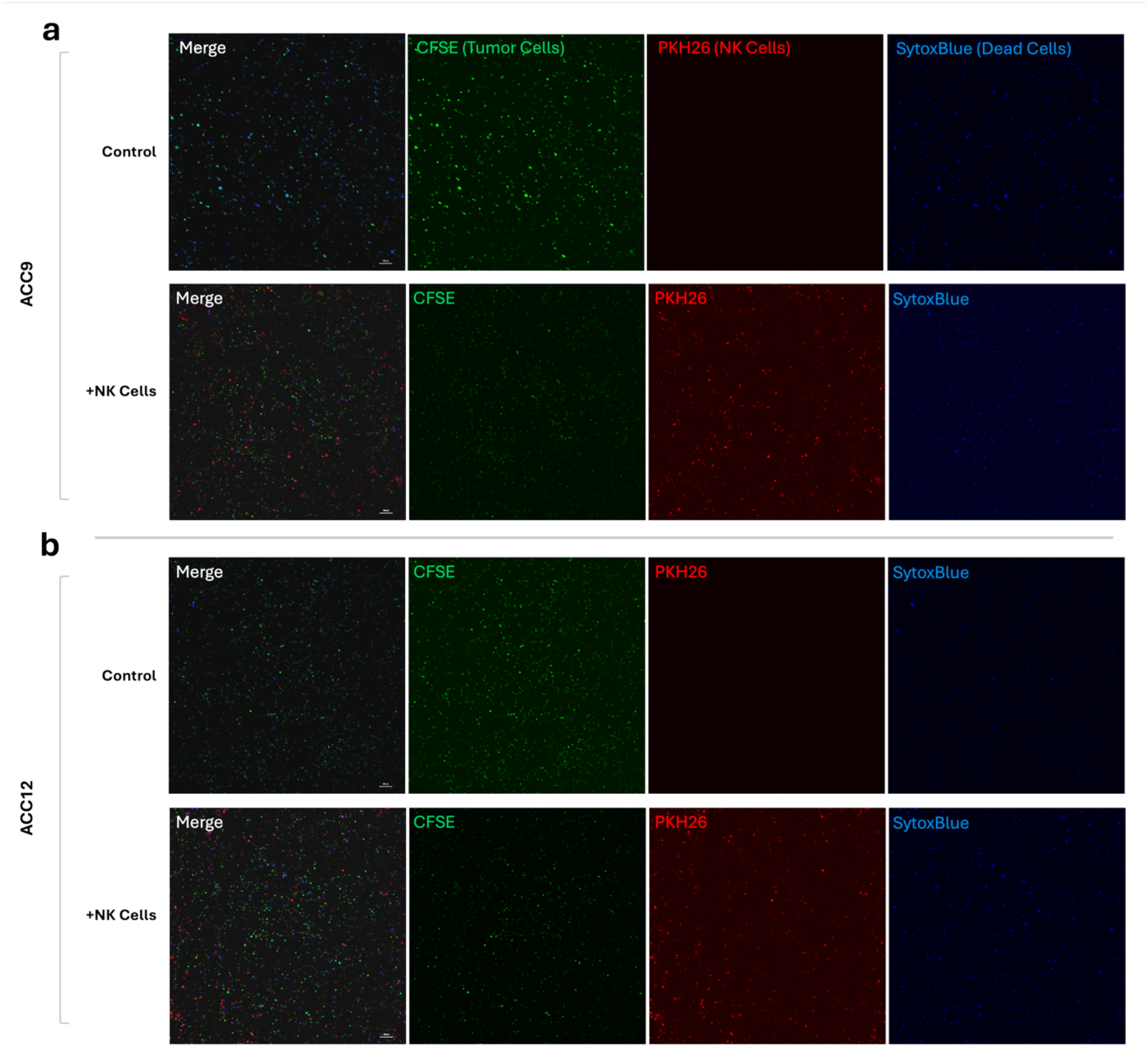
NK cell-based immunotherapy in PTCs. In a) ACC9 and b) ACC12 PTCs, addition of NK-92 cells results in increased NK-92 cell infiltration. CFSE – prelabeled tumor cells; PKH26, prelabeled NK-92 cells; SYTOX Blue – Dead cells.

## 3. Discussion

In these studies, we demonstrated the capability to generate 3D tumor models from clinical ACC biospecimens – a platform that has been challenging to establish for this rare malignancy. After establishment, ACC PTCs showed marker expression similar to primary ACC. Additionally, PTCs maintained proliferation over time, and produced cortisol. Transcriptomic analysis confirmed that PTCs resembled their matching tumors, establishing biological fidelity of the model.

We then evaluated ACC PTCs in chemotherapy screening studies utilizing clinically relevant drug compounds, mitotane and EDP. Despite limited clinical efficacy, we did observe partial cytotoxic effects, which is consistent with the modest clinical efficacy of these agents. To identify alternative treatments, we screened a panel of target therapies: Wnt, IGF-2, and ATR inhibitors, as well as a p53 activator. We screened these compounds in NCI-H295R cells to establish dose-response curves. ATR inhibitors (VX-803 and elimusertib) demonstrated the lowest effective concentrations, followed by the Wnt inhibitor, tegavivint. The Wnt inhibitor XAV-939 showed minimal response even at high concentrations, while the p53 activator, NSC59984 only reduced viability at high concentrations.

Based on established dysregulation of Wnt and IGF-2 signaling in ACC, we evaluated combinatorial treatment with tegavivint and linsitinib. In NCI-H295R cells, specific drug ratios produced additive effects (**Supplemental Figure 1e**). However, when tested in ACC PTC, both drugs were effective as monotherapy but combination treatment showed no additive benefit. This discrepancy between NCI-H295R and patient-derived models underscores the importance of using primary tumor platforms for therapeutic screening.

Finally, we evaluated cellular immunotherapy approaches, which have not been extensively evaluated in ACC patient derived models. We tested NK-92 natural killer cells and TALL-104 CD8+ T cells in NCI-H295R constructs. TALL-104 cells, despite demonstrating efficacy in other tumor models, showed limited cytotoxicity against ACC. In contrast, NK-92 cells effectively infiltrated and killed ACC cells in both NCI-H295R constructs and PTCs, demonstrating consistent NK cell-mediated cytotoxicity across ACC models. This differential response may reflect the immunosuppressive effects of cortisol, which ACC tumors frequently produce in excess. Previous studies have shown that glucocorticoid excess depletes T cell populations and is associated with resistance to T cell-based checkpoint inhibitor therapies in ACC. In contrast, NK cells function through innate immunity mechanisms that do not require antigen presentation, potentially making them less susceptible to cortisol-mediated immunosuppression.

Our PTC platform employs a defined, bioengineered hyaluronic acid and collagen hydrogel that eliminates the batch-to-batch variability inherent to Matrigel and other basement membrane extracts derived from murine sarcomas.^29^ This reproducibility comes at a tradeoff: the defined system lacks endogenous cytokines and growth factors, limiting long-term primary cell culture without supplementation. Despite these constraints, our ACC PTCs demonstrated faithful recapitulation of patient tumor biology, including cortisol production, marker expression, and transcriptomic signatures. The platform enabled systematic evaluation of both conventional and targeted therapeutics, revealing ATR inhibition as a promising strategy and establishing proof-of-concept for NK cell-based immunotherapy. These patient-derived models provide a scalable, reproducible system for therapeutic discovery in ACC—a rare malignancy where preclinical tools have historically been scarce. By bridging the gap between cell line studies and clinical translation, ACC PTCs offer a path forward for identifying and validating novel treatments for this lethal disease.

## 4. Materials and Methods

### 4.1. Cell line cultures

NCI-H295R, NK-92, and TALL-104 cells were obtained from American Type Culture Collection (ATCC). H295R cells were cultured in DMEM-F12 (ATCC) supplemented with 2.5% Nu-serum (Corning) and 1X ITS+ Premix Universal Culture Supplement (Corning). Cells were harvested at 80% confluence and cultured for less than 8 passages. NK-92 cells were expanded in MEM ɑ supplemented with 0.1 mM β-mercaptoethanol, 0.02 mM Folic Acid, 0.2 mM myo-Inositol, 125 IU/ul human recombinant Interleukin-2 (IL-2), and 12.5% each of fetal bovine serum (FBS) and horse serum. TALL-104 cells were cultured in IMDM with 0.5 ug/mL D-mannitol, 0.1 mM β-mercaptoethanol, 2.5 ug/mL human serum albumin (HSA), 100 IU/mL IL-2, and 20% FBS.

### 4.2. ACC biospecimen acquisition and processing

Resected tumor samples were transferred to the laboratory on RPMI-1640 for processing on the same day. Samples were washed in 2% penicillin-streptomycin in PBS 3 times for 5 minutes followed by 2 washes of 2% penicillin-streptomycin in DMEM-F12. Tissues were finely dissociated with the use of a scalpel and placed into DMEM-F12 supplemented with 10% collagenase/hyaluronidase (StemCell), 22,000 units/mL protease (VitaCyte), and 100,000 units/mL collagenase HA (VitaCyte) for less than 2 hours at 37 °C with gentle shaking. Upon sufficient digestion, the tissue mix was neutralized with DMEM supplemented with 10% FBS and passed through a 100 μm pore filter, to be centrifuged. Isolated pellets were incubated with 1X red blood cell lysis buffer, recentrifuged, counted and assayed for viability. Cells were then expanded in Matrigel coated 6-well plates with DMEM:F12 supplemented with 10 nM HEPES, 1x Glutamax, 1x B27, 1.25 mM N-acetylcysteine, 10mM nicotinamide, 0.5 uM A-83-01, 10nM [Leu15]-Gastrin I, 50 ng/mL EGF, 0.5 ug/mL R-spondin 1, 0.1 ug/mL FGF10, 30 ng/mL Wnt3a, 0.1 ug/mL Noggin, and 1 uM PGE2.

### 4.3. ACC tumor construct biofabrication

Thyolated hyaluronic acid and heparin (Advanced Biomatrix) was reconstituted to a 1% w/v concentration with 0.01% w/v Irgacure in ultrapure water. Methacrylated type 1 collagen (Advanced Biomatrix) was reconstituted with 20 mM acetic acid to a final concentration of 6 mg/mL. H295R or patient tumor derived cells were resuspended in a hydrogel mix of 3 parts neutralized collagen to 1-part hyaluronic acid at a concentration of 10,000 cells/ul. Constructs of 10 ul were dispensed onto PDMS-coated 48-well plates and crosslinked for 2 seconds with a 365 nm UV probe (BlueWave RediCure 365, Dymax, Torrington, CT).

### 4.4. Proliferation and viability assays

ACC PTC cell proliferation and viability were measured using ATP quantification and LIVE/DEAD staining paired with fluorescent microscopy. Cell viability via ATP quantification was carried out using the Promega® CellTiter-Glo® 3D - Superior Cell Viability Assay according to the manufacturer’s instructions. LIVE/DEAD-staining was carried out using the Viability/Cytotoxicity kit for mammalian cells (Invitrogen) according to the manufacturer’s instructions. Fluorescent microscopy imaging of calcein AM-stained viable cells and ethidium homodimer-1-stained dead cell nuclei was performed using Nikon A1R and AXR confocal microscopes.

### 4.5. Cortisol measurements

During PTC culture, media aliquots were reserved on days 4 and 7 and frozen until cortisol was quantified. Specifically, cortisol concentrations were determined by human Cortisol ELISA kit (cat. no. ab108665; Abcam), following the manufacturer’s instructions.

### 4.6. Immunofluorescent staining and microscopy

Immunofluorescence (IF) was performed on sectioned paraffin-embedded tumor constructs or fixed cells cultured for 3 to 7 days on X-tra® slides to analyze traditional ACC biomarker presence of SF-1 (ab240394, Abcam), Melan A (ab210546, Abcam), inhibin-a (ab47720, Abcam), IGF-2 (ab226989, Abcam), and b-catenin (ab ab223075, Abcam). Nuclei were stained with DAPI (P36962, ThermoFisher). The slides were imaged with Nikon AXR confocal microscope at 40X magnification at room temperature.

### 4.7. Bulk RNA-sequencing

Total RNA was isolated from PTCs, their matching tumors, and the NCI-H295R cell line. Samples ACC 1 and ACC 4 are paired and originate from the primary and metastatic lesion of the same patient, respectively. Sequencing libraries were prepped using the SMARTer® Stranded Total RNA-Seq - Pico Input Mammalian Kit v3 (Takara bio, cat#634485) and sequenced on an Illumina NovaSeq6000 to generate paired-end sequencing reads. Salmon¹ v1.3.0 was used to align the reads to the GRCh38 human reference genome and to quantify the transcripts. Resulting data were preprocessed using DESeq2² v1.44.0 for R. PCA plots were created using PCAtools³ v2.16.0 for R. Normalized expression for the overlap of the top 50 PC 1 and PC 2 loadings was plotted on a heatmap using pheatmap^4^ v1.0.12 for R.^30,31^

### 4.8. Mitotane and EDP drug studies

ACC PTCs were treated with mitotane (S1732, Selleckchem, Houston, TX), etoposide (S1225, Selleckchem), doxorubicin (S1208, Selleckchem), and cisplatin (S1126, Selleckchem). The tumor constructs or 2D cell cultures were exposed to clinically relevant drug conditions of mitotane (8 µg/mL or 20 µg/mL), a combination of etoposide, doxorubicin, and cisplatin (EDP) at concentrations of 0.25 µg/mL, 0.1 µg/mL D, 0.1 µg/mL, respectively, or at concentrations of 25 µg/mL, 10 µg/mL D, 10 µg/mL, respectively. In two drug conditions, mitotane at 8 µg/mL was combined with EDP at 0.25 µg/mL, 0.1 µg/mL D, 0.1µg/mL or mitotane at 20 µg/mL and EDP at 25µg/mL, 10 µg/mL D, 10 µg/mL respectively. After 3 days media was replaced with fresh media containing the aforementioned drug concentrations. Viability was assessed after 72 hours using ATP quantification and LIVE/DEAD staining as described above.

### 4.9. Wnt, IGF2, P53, and ATR pathway inhibition studies

NCI-H295R constructs were fabricated as above and fresh media was added to constructs with varying concentration of Linsitinib, Tegatrabetan, XAV-939, VX-803, Elimusertib, and NSC 59984. Media and chemotherapeutic compounds were refreshed on day 4 of culture, and samples were stained on days 1, 4, and 7 with calcein-AM and ethidium homodimer-1, live and dead cell indicators (Thermo Fisher). Constructs were imaged using a Nikon AxR confocal microscope with a 10x objective.

ACC PTCs were fabricated as above and treated with concentrations increasing logarithmically between 0 and 1×10^5^ nM (n=5 per concentration) of the above compounds. Media and chemotherapeutic agents were replenished on day 3 of cultured and cell metabolic activity was assayed with CellTiter-Glo Luminescent Cell Viability Assay (Promega). IGF-R1 and Wnt/β-Catenin combinatorial pathway inhibition was carried out by initially treating constructs with Linsitinib and Tegatrabetan separately to determine IC50 values. Next samples were set to the determined IC50 values of one compound while varying the concentration of the other in combinatorial treatment. Data was plotted into Prism and dose response curves were fitted into values.

### 4.10. Cellular immunotherapy studies

H295R or patient derived cells were stained with carboxyfluorescein succinimidyl ester (CFSE) (Thermo Fisher) and embedded at a concentration of 5,000 cells/ul into a hydrogel consisting of 2% w/v methacrylated gelatin (Advanced Biomatrix), 4 mM 10 kDa thyol-reactive polyethylene glycol diacrylate (Creative PEGWorks), 1 mg/mL rat tail collagen type I (Advanced Biomatrix), and 0.1% w/v Irgacure. Constructs were pipetted as 5 ul aliquots into PDMS-coated 48-well plates, UV-crosslinked as above, and incubated at 37 °C for 20 minutes. Constructs were allowed to incubate overnight following the addition of complete media. TALL-104 or NK-92 cells stained with PKH26 (Sigma) were then added for a coculture ratio of 4:1, lymphocytes to cancer cells. After 3 days of co-culture, constructs were stained with SYTOX blue (Thermo Fisher) and imaged using the confocal microscope.

## Acknowledgements

PD and AS acknowledge funding through the Ohio State University Comprehensive Cancer Center’s Cancer Biology Program Seed Award Program and NIH grant R21CA277083. AS acknowledge awards from the Pelotonia foundation. PD acknowledges funding from the American Association of Endocrine Surgeons and the Society of University Surgeons. We also acknowledge resources from the Campus Microscopy and Imaging Facility (CMIF) and the OSU Comprehensive Cancer Center (OSUCCC) Microscopy Shared Resource (MSR), The Ohio State University. This facility is supported in part by NIH/NCI grant P30 CA016058.

## Supplementary Materials

**Supplemental Figure 1:**
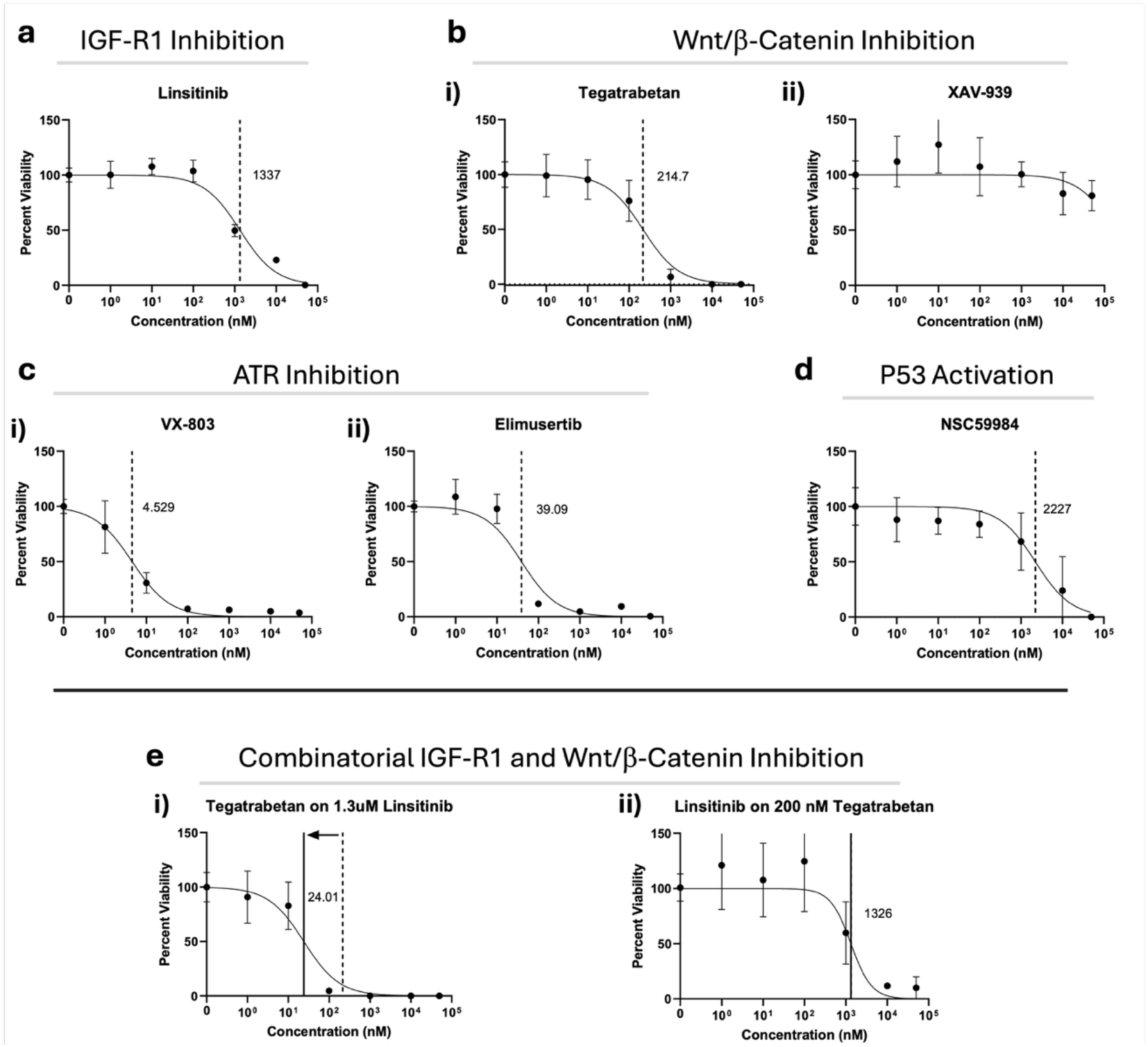
Cell line-based drug studies. Generation of curves for a) IGF-R1 inhibitor linsitinib, b) Wnt/β-Catenin inhibitors i) tegatrabetan, and ii) XAV-939, c) ATR inhibitors i) VX-803 and ii) elimusertib, d) P53 activator NSC59984, and e) combinatorial therapy with tegatrabetan and linsitinib. Arrows between vertical dashed lines show changes in IC50 during combinational treatments

**Supplemental Figure 2:**
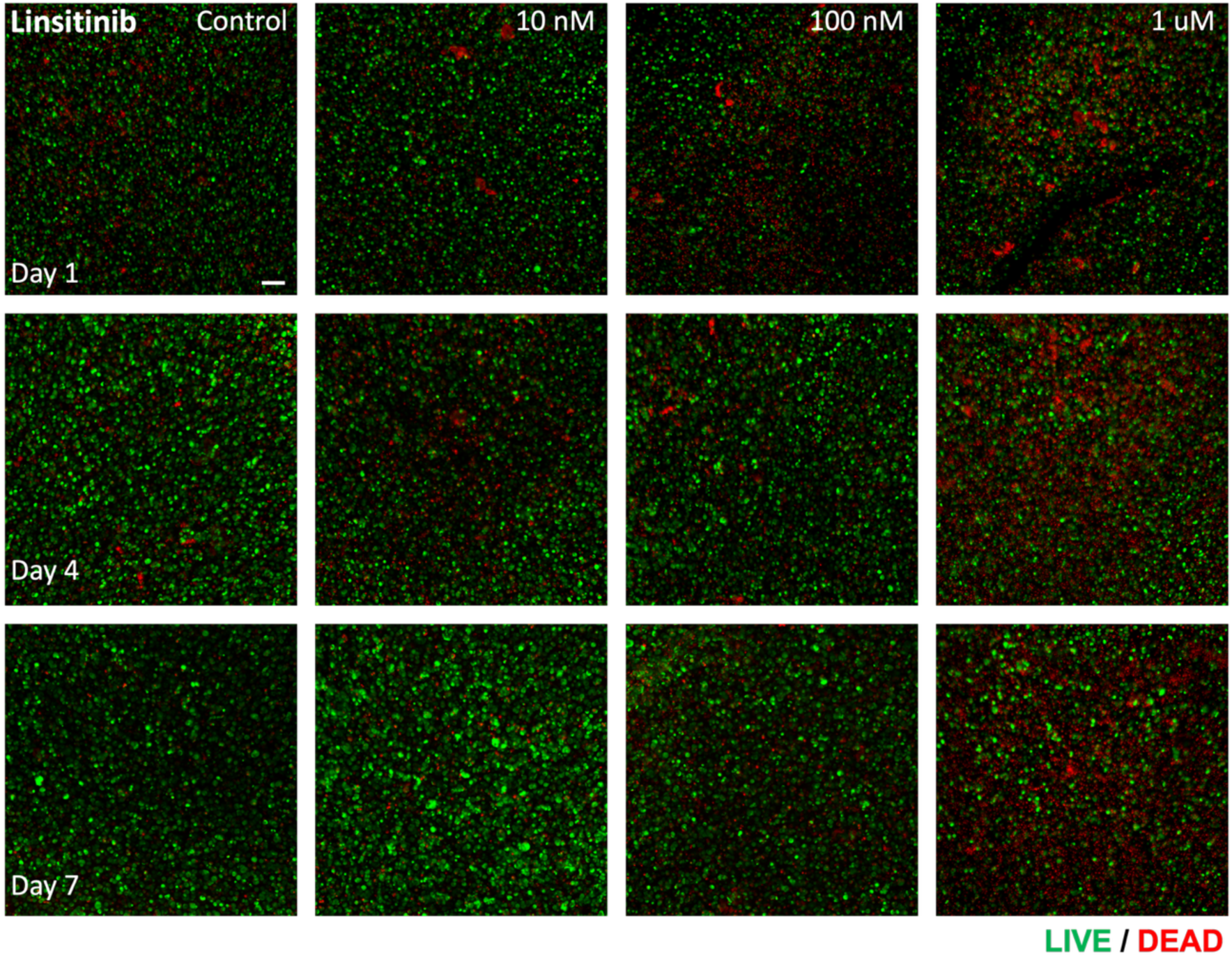
LIVE/DEAD viability panel for cell line-based ACC constructs treated with linsitinib. LIVE/DEAD staining: Green – Calcein AM-stained viable cells; Red – Ethidium homodimer-1-stained dead cell nuclei. Scale bar = 100 um.

**Supplemental Figure 3:**
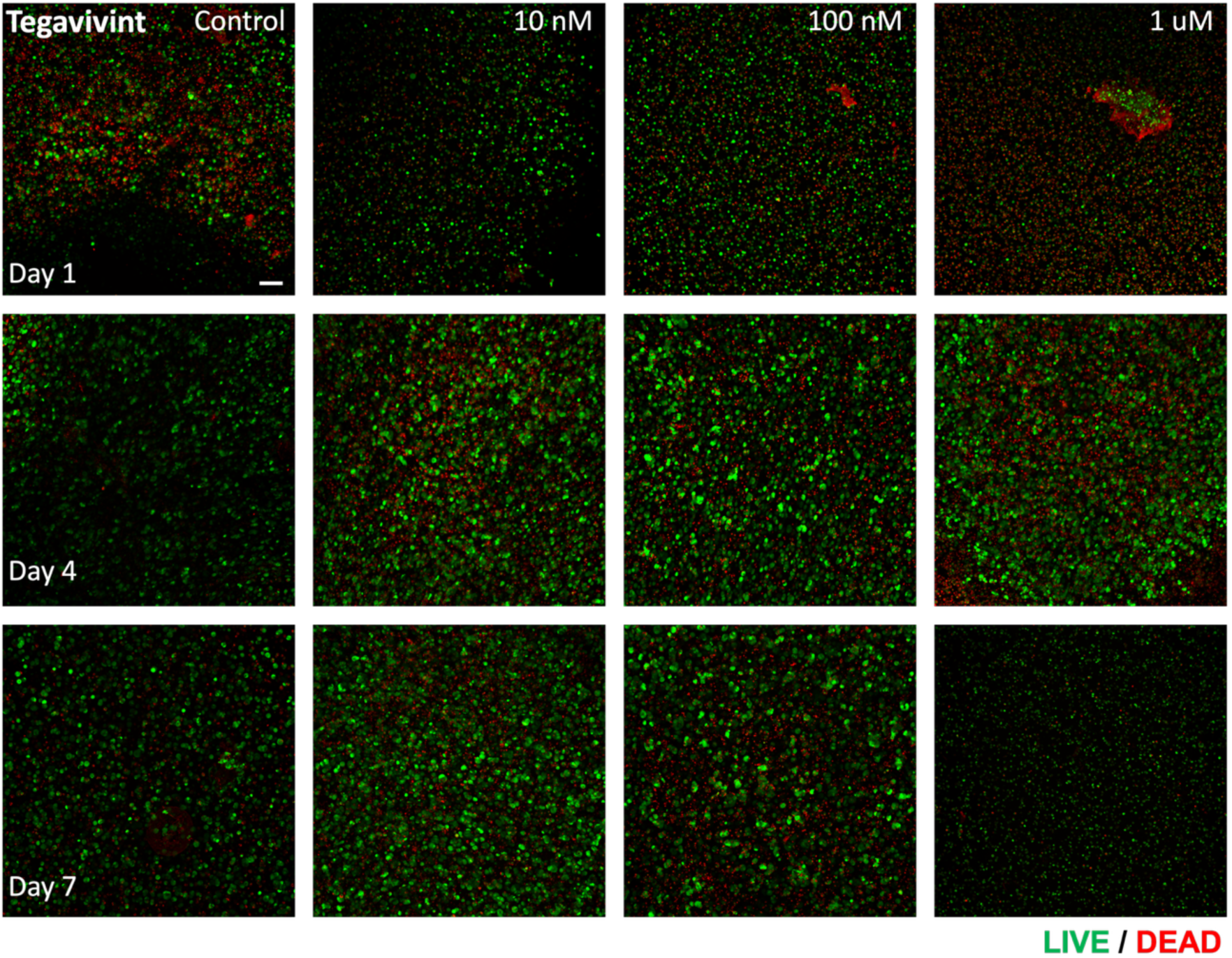
LIVE/DEAD viability panel for NCI-H295R constructs treated with tegavivint (tegatrabetan). LIVE/DEAD staining: Green – Calcein AM-stained viable cells; Red – Ethidium homodimer-1-stained dead cell nuclei. Scale bar = 100 um.

**Supplemental Figure 4:**
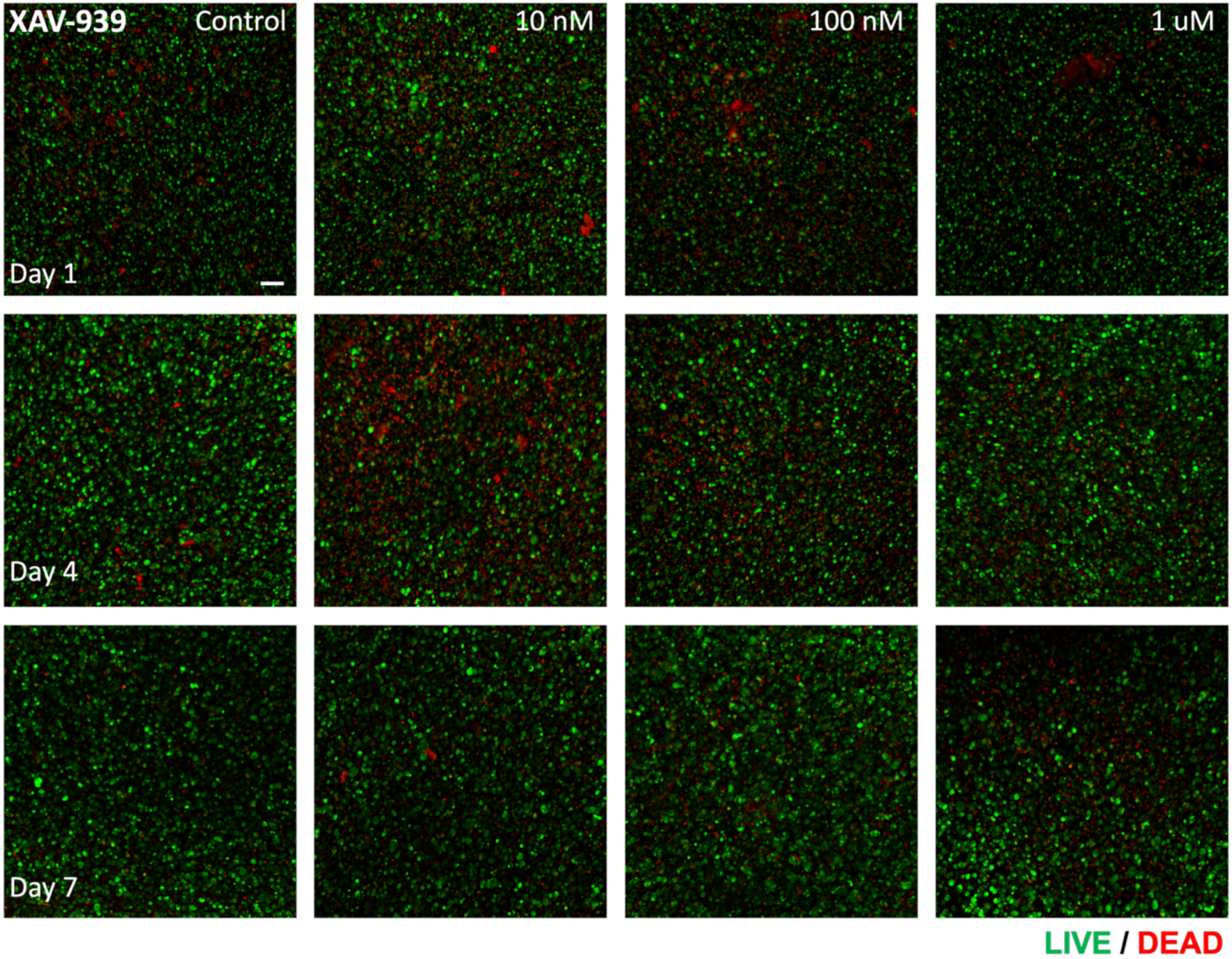
LIVE/DEAD viability panel for NCI-H295R constructs treated with XAV-939. LIVE/DEAD staining: Green – Calcein AM-stained viable cells; Red – Ethidium homodimer-1-stained dead cell nuclei. Scale bar = 100 um.

**Supplemental Figure 5:**
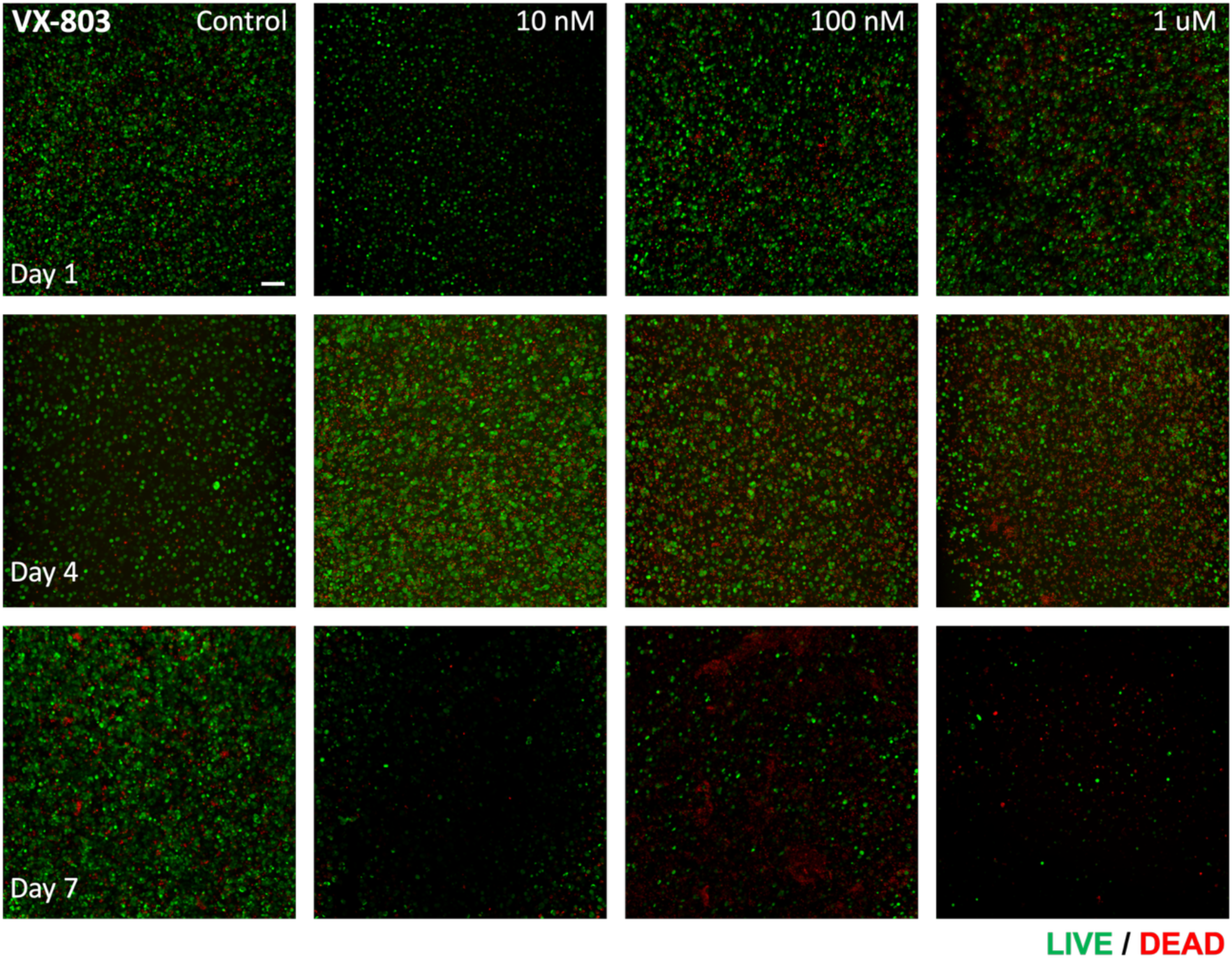
LIVE/DEAD viability panel for NCI-H295R constructs treated with VX-803. LIVE/DEAD staining: Green – Calcein AM-stained viable cells; Red – Ethidium homodimer-1-stained dead cell nuclei. Scale bar = 100 um.

**Supplemental Figure 6:**
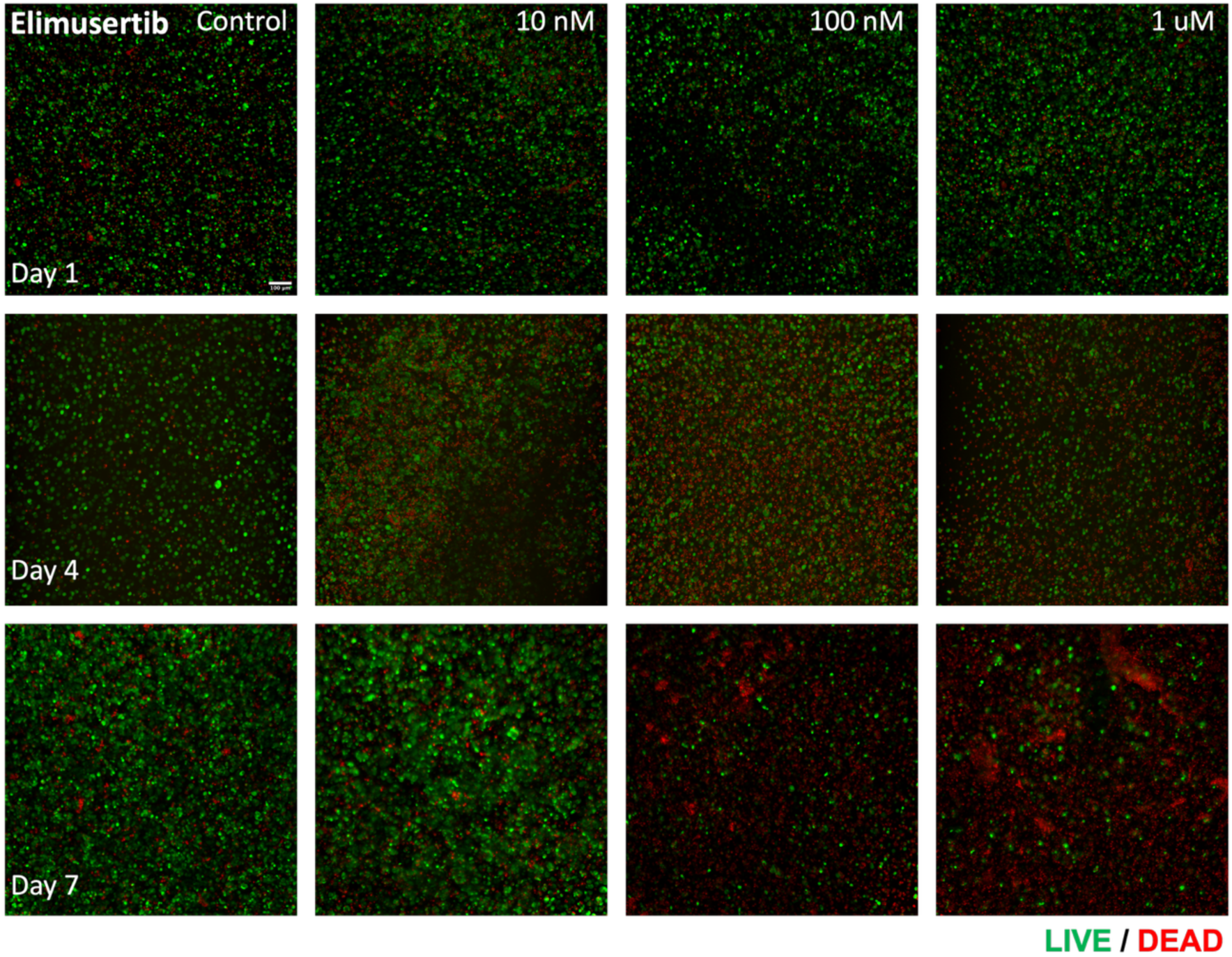
LIVE/DEAD viability panel for NCI-H295R constructs treated with elimusertib. LIVE/DEAD staining: Green – Calcein AM-stained viable cells; Red – Ethidium homodimer-1-stained dead cell nuclei. Scale bar = 100 um.

**Supplemental Figure 7:**
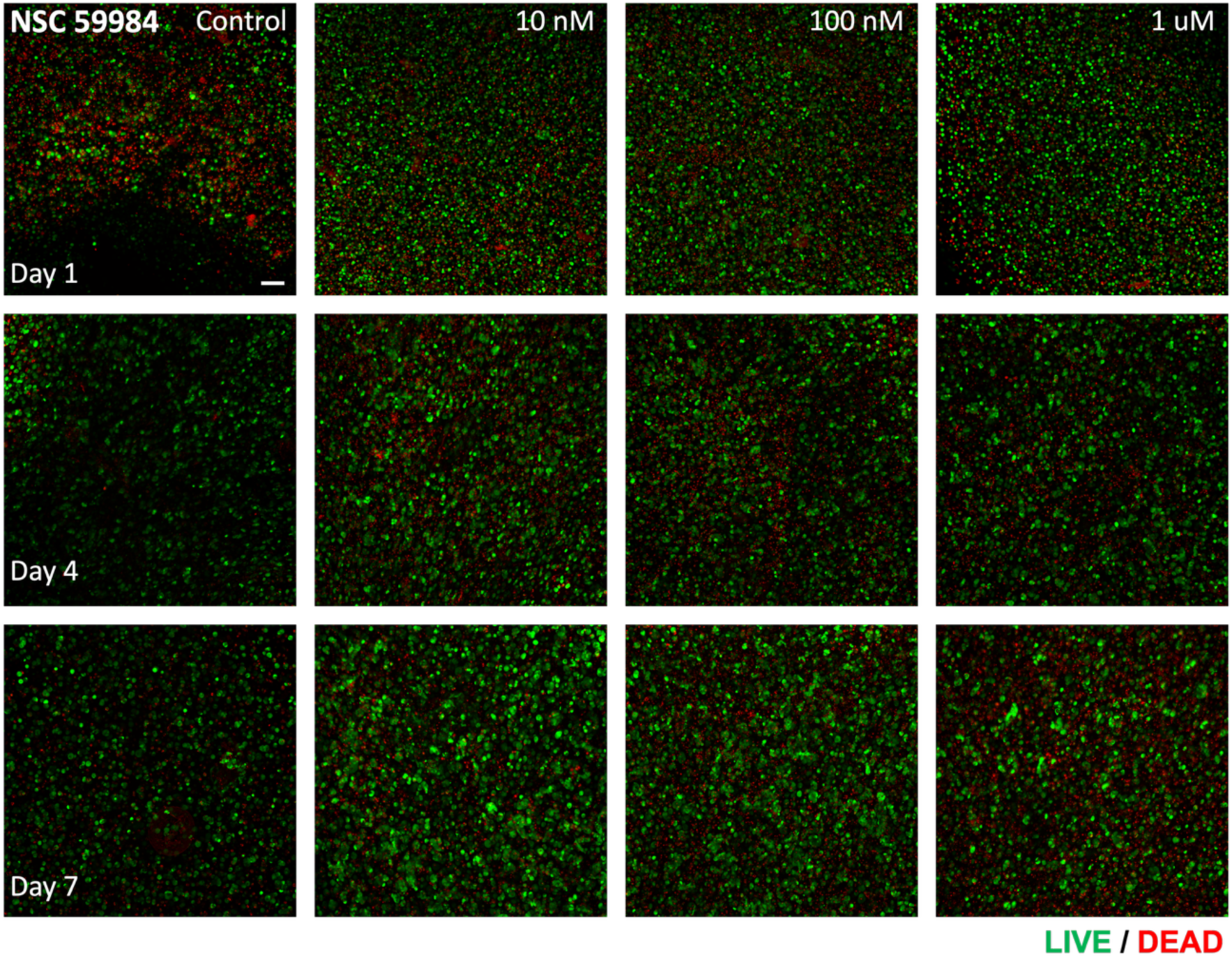
LIVE/DEAD viability panel for NCI-H295R constructs treated with NSC 59984. LIVE/DEAD staining: Green – Calcein AM-stained viable cells; Red – Ethidium homodimer-1-stained dead cell nuclei. Scale bar = 100 um.

**Supplemental Figure 8:**
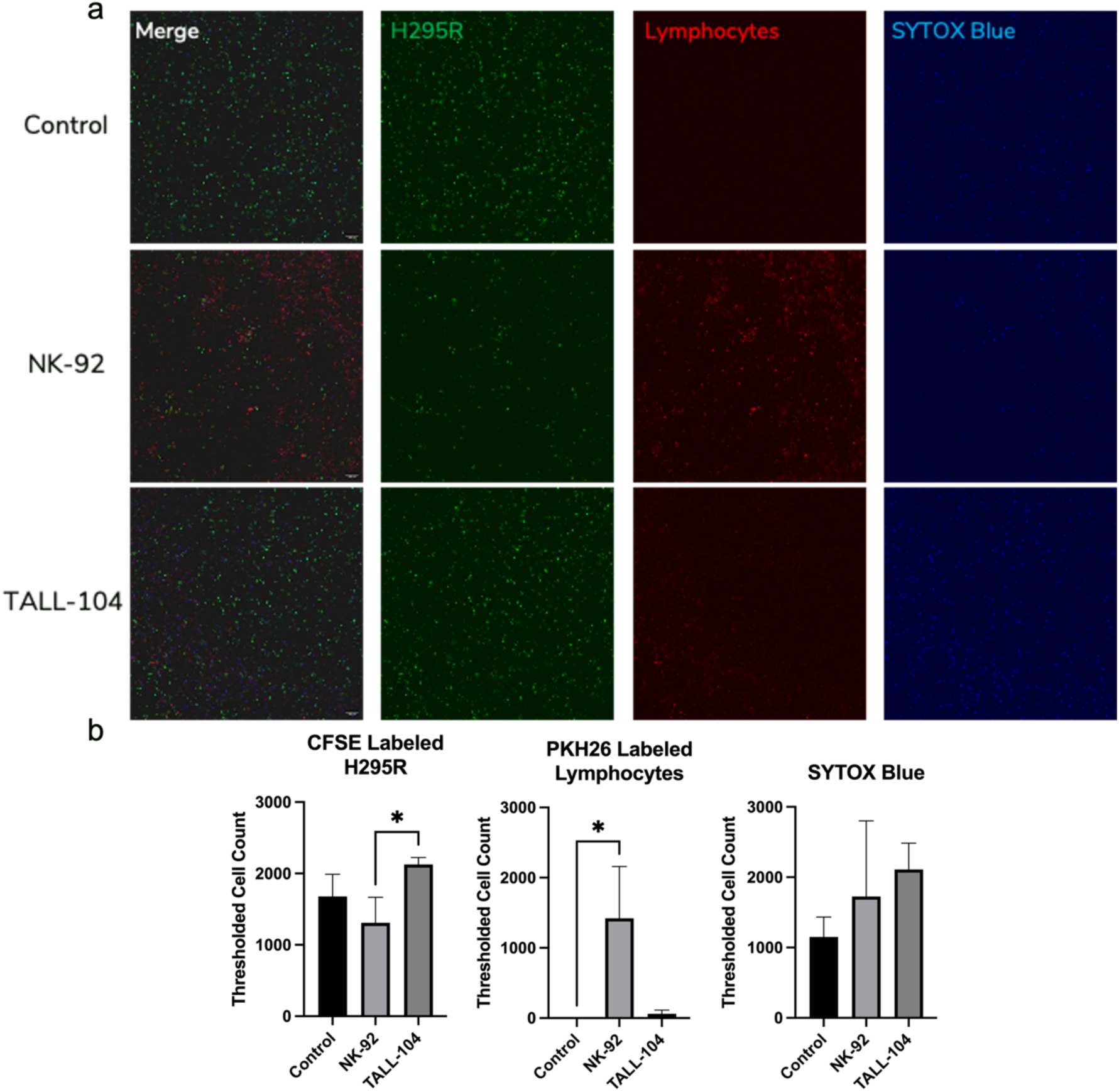
NK-92 and TALL-104 effect on NCI-H295R constructs. a) Confocal images displaying addition of NK-92 and TALL-104 cells to H295R constructs. b) Quantification of fluorescent signal for CFSE, PKH26, and SYTOX Blue. Addition of NK-92 cells results in decreased tumor cell presence, increased NK-92 cell infiltration, and increased dead cell numbers, while addition of TALL-104 cells observes overall less infiltration. CFSE – prelabeled tumor cells; PKH26, prelabeled NK-92 and TALL cells; SYTOX Blue – Dead cells.

